# Regulation of translation by lysine acetylation in *Escherichia coli*

**DOI:** 10.1101/2022.05.02.490376

**Authors:** Sarah C. Feid, Hanna E. Walukiewicz, Xiaoyi Wang, Ernesto S. Nakayasu, Christopher V. Rao, Alan J. Wolfe

## Abstract

N^*ε*^-lysine acetylation is a common post-translational modification observed in diverse species of bacteria. Aside from a few central metabolic enzymes and transcription factors, little is known about how this post-translational modification regulates protein activity. In this work, we investigated how lysine acetylation affects translation in *Escherichia coli*. In multiple species of bacteria, ribosomal proteins are highly acetylated at conserved lysine residues, suggesting that this modification may regulate translation. In support of this hypothesis, we found that the addition of the acetyl donors, acetyl phosphate or acetyl-Coenzyme A, inhibits translation but not transcription using an *E. coli* cell-free system. Further investigations using *in vivo* assays revealed that acetylation does not appear to alter the rate of translation elongation but rather increases the proportion of dissociated 30S and 50S ribosomes, based on polysome profiles of mutants or growth conditions known to promote lysine acetylation. Furthermore, ribosomal proteins are more acetylated in the disassociated 30S and 50S ribosomal subunit than in the fully assembled 70S complex. The effect of acetylation is also growth rate dependent, with disassociation of the subunits most pronounced during late exponential and early stationary phase growth – the same growth phase where protein acetylation is greatest. Collectively, our data demonstrate that lysine acetylation inhibits translation, most likely by interfering with subunit association. These results have also uncovered a new mechanism for coupling translation to the metabolic state of the cell.

**IMPORTANCE:** Numerous cellular processes are regulated in response to the metabolic state of the cell. One such regulatory mechanism involves lysine acetylation, a covalent modification involving the transfer of an acetyl group from the central metabolites acetyl coenzyme A or acetyl phosphate to a lysine residue in a protein. This post-translational modification is known to regulate some central metabolic enzymes and transcription factors in bacteria, though a comprehensive understanding of its effect on cellular physiology is still lacking. In the present study, lysine acetylation was also found to inhibit translation in *Escherichia coli* by impeding ribosome association, most likely by disrupting salt-bridges along the binding interface of the 30S and 50S ribosomal subunits. These results further our understanding of lysine acetylation by uncovering a new target of regulation, protein synthesis, and aid in the design of bacteria for biotechnology applications where the growth conditions are known to promote lysine acetylation.

## INTRODUCTION

N^ε^-lysine acetylation is a post-translational modification found in all domains of life, and it is consistently observed in diverse bacterial species (1–3). This modification neutralizes the positive charge of lysine residues by covalently attaching an acetyl group to the amino group of the lysine side chain. While some acetylated lysines are known to alter protein activity, the vast majority remain uncharacterized (4, 5). One under-investigated target of lysine acetylation is the bacterial ribosome, whose proteins are consistently acetylated at conserved sites in diverse bacteria (**Fig. 1**) (6, 7). Despite being a common target of acetylation, little is known about the effect of acetylation on the ribosome and translation in general.

**Figure 1.**
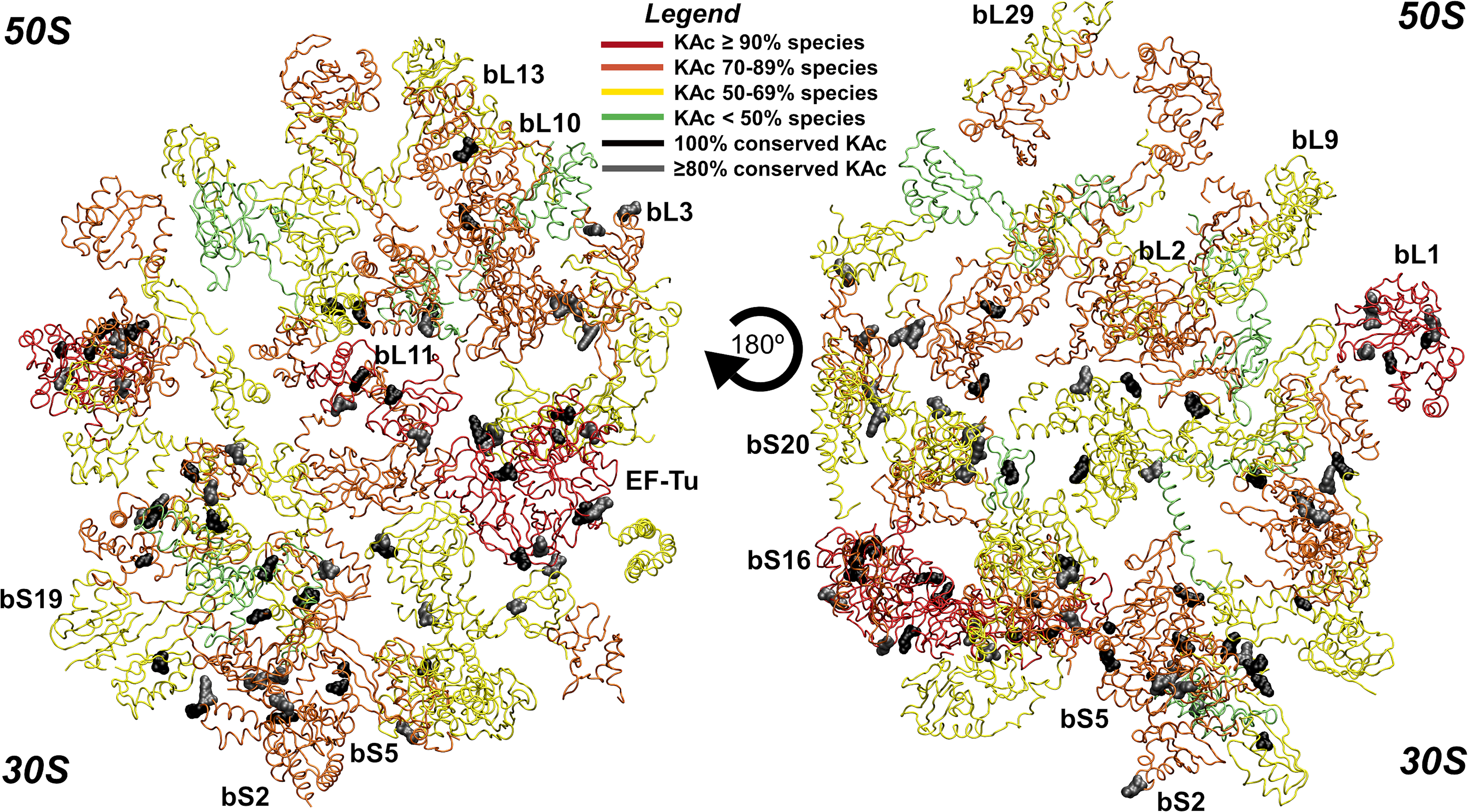
Conserved lysine acetylation sites on bacterial ribosomal proteins. Acetylation sites were extracted from a previously published proteomics dataset of 48 phylogenetically distant bacteria (6). The degree of acetylation across different species and conserved sites were mapped onto a ribosomal protein structure deposited in PDB (accession number 5UYK) using Visual Molecular Dynamics software v.1.9.3. Polypeptide chains are colored according to the percentage of species that each ribosomal protein is acetylated, while the RNA molecules were hidden. Acetylation on sites conserved that were invariant or conserved in at least 80% of the 48 bacterial species are highlighted in black and grey, respectively.

Lysines can be acetylated by two distinct mechanisms: enzymatically by lysine acetyltransferases using acetyl coenzyme A (acetyl-CoA) as the acetyl donor and non- enzymatically using acetyl phosphate or, more rarely, acetyl-CoA, as the donor (8, 9). In *Escherichia coli*, *Neisseria gonorrhoeae* and *Bacillus subtilis,* the majority of acetylations occur non-enzymatically (10–13). Some acetylations are removed by lysine deacetylases (14–16). Most bacteria express only one or two lysine deacetylases, which do not appear to act upon the majority of acetyllysines. Thus, most lysine acetylations are believed to be not reversed (17).

Proteins are principally acetylated when cells enter stationary phase during growth on excess carbon (7, 18, 19). To maintain flux through glycolysis, *E. coli* can ferment excess carbon to acetate through the Pta-AckA pathway, where phosphotransacetylase (Pta) converts acetyl-CoA and inorganic phosphate to acetyl phosphate and free CoA, and then acetate kinase (AckA) converts acetyl phosphate and ADP to acetate and ATP (20). More flux through this pathway increases the intracellular concentrations of acetyl phosphate, which is directly tied to the level of non-enzymatic acetylation in the cell (10, 13).

As cells enter stationary phase, the ribosomes undergo several changes. The rate of protein synthesis decreases as does the rate of translation elongation (21). The number of 70S ribosomes decreases, either due to subunit dissociation or formation of 100S ribosomes, and the remaining 70S become less active (22). Ribosome population differences that depend on the limiting nutrient suggests that metabolism regulates ribosome function; for example, phosphorus-limited *E. coli* maintain the same growth rate and protein levels as carbon- or nitrogen-limited *E. coli* but with fewer ribosomes (23). In support of a mechanism whereby acetylation regulates ribosome activity, recent work suggests that an accumulation of acetylations during stationary phase decreases the rate of elongation (24).

In this work, we investigated the effect of lysine acetylation on translation in *E. coli*. Using an *in vitro* transcription/translation assay, we found that acetyl donors inhibit translation but not transcription. To better understand the mechanism, we performed polysome profiling and found that fewer ribosomes form 70S complexes in high acetylation mutants and/or under growth conditions known to promote acetylation. In contrast, under these conditions, we did not observe an acetylation-dependent effect on elongation rate. These results suggest that lysine acetylation inhibits translation by promoting disassociation or inhibiting association of the ribosome.

## Results

### Acetyl donors inhibit translation

Ribosomal proteins are highly acetylated, and the acetylated lysine residues are highly conserved in diverse species of bacteria (**Fig. 1**). Therefore, we hypothesized that ribosome acetylation would affect translation. To test this hypothesis, we used a cell-free transcription/translation system derived from *E. coli* cell lysates to measure production of the green fluorescent protein (deGFP, a variant of eGFP optimized for cell-free synthesis) from a σ^70^-dependent promoter on a plasmid in the presence and absence of the acetyl donors, acetyl-CoA or acetyl phosphate (**Fig. 2A**) (25). The addition of either acetyl donor, at the upper range of their physiologically relevant concentrations, strongly inhibited the production of deGFP as determined by fluorescence. While acetyl-CoA can non-enzymatically acetylate lysine residues, its contribution to *in vivo* acetylation is difficult to determine as it essential in most organisms (7). Going forward, we chose to focus on acetyl phosphate, as it has been identified as the primary non-enzymatic acetyl donor in several bacteria (10–13). The concentration of acetyl phosphate varies based on the growth conditions and, in *E. coli*, can reach 5 mM (26). Using a spread of physiologically relevant acetyl phosphate concentrations, we found that deGFP production was inhibited in a dose-dependent manner (**Fig. 2B**). To determine whether the addition of acetyl phosphate was inhibiting transcription or translation, we used quantitative PCR to measure relative *degfp* mRNA levels. Consistent with a mechanism whereby acetyl phosphate inhibits translation, we found that mRNA levels did not decrease with increasing concentrations; at lower acetyl phosphate concentrations, mRNA increased relative to the untreated control. (**Fig. 2C**). These results suggest acetyl donors inhibit translation, likely by acetylating ribosomal proteins.

**Figure 2.**
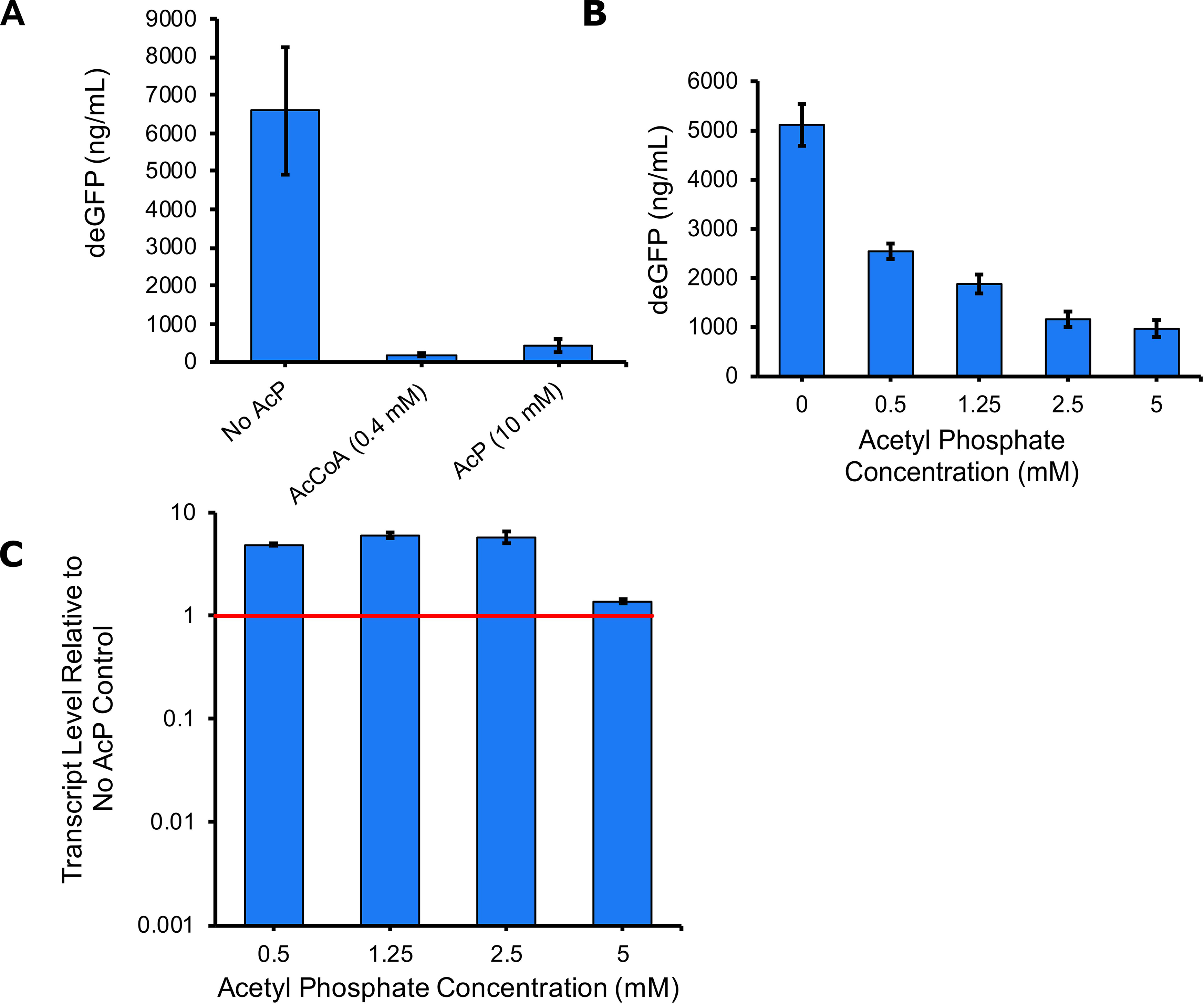
Addition of acetyl donors inhibits translation but not transcription. deGFP synthesis by a cell-free transcription translation system was measured in the presence of **(A)** acetyl-CoA, or acetyl phosphate and **(B)** various concentrations of acetyl phosphate**. (C)** RNA was isolated from reactions for qRT-PCR. Expression of deGFP transcript was determined relative to the no acetyl phosphate control. Error bars represent the standard deviation from two replicates.

### Conditions promoting acetylation do not affect elongation

Previous work suggested that acetylation reduces the elongation rate (24). Such a mechanism could potentially explain the decreased rates of translation observed using the *in vitro* cell-free system. To test this hypothesis, we measured elongation rates using a LacZ induction assay (21, 27). To manipulate acetylation levels, or more precisely acetyl phosphate concentrations, we grew a Δ*pta* mutant in MOPS glucose minimal medium with or without acetate. This allowed us to manipulate the direction of the Pta-AckA pathway. When grown on MOPS glucose, this mutant does not produce acetyl phosphate as phosphate acetyltransferase (Pta) is needed to convert acetyl-CoA to acetyl phosphate and, thus, acetylation is low (20). When the growth medium is supplemented with sodium acetate, the Δ*pta* mutant accumulates acetyl phosphate as acetate kinase (AckA) assimilates acetate to acetyl phosphate; thus, acetylation is high (20). When *lacZ* was induced in stationary phase, we observed no difference in the elongation rate between the Δ*pta* mutant grown on 0.2% glucose versus 0.2% glucose supplemented with 0.27% sodium acetate (**Fig. 3**). The slight, non-significant decrease in elongation rate for the Δ*pta* mutant in both conditions is attributed to a modest growth defect of Δ*pta* relative to wild type (**Fig. S1**). These results suggest that acetyl donors do not alter the rate of translation elongation, at least under the conditions tested in these experiments.

**Figure 3.**
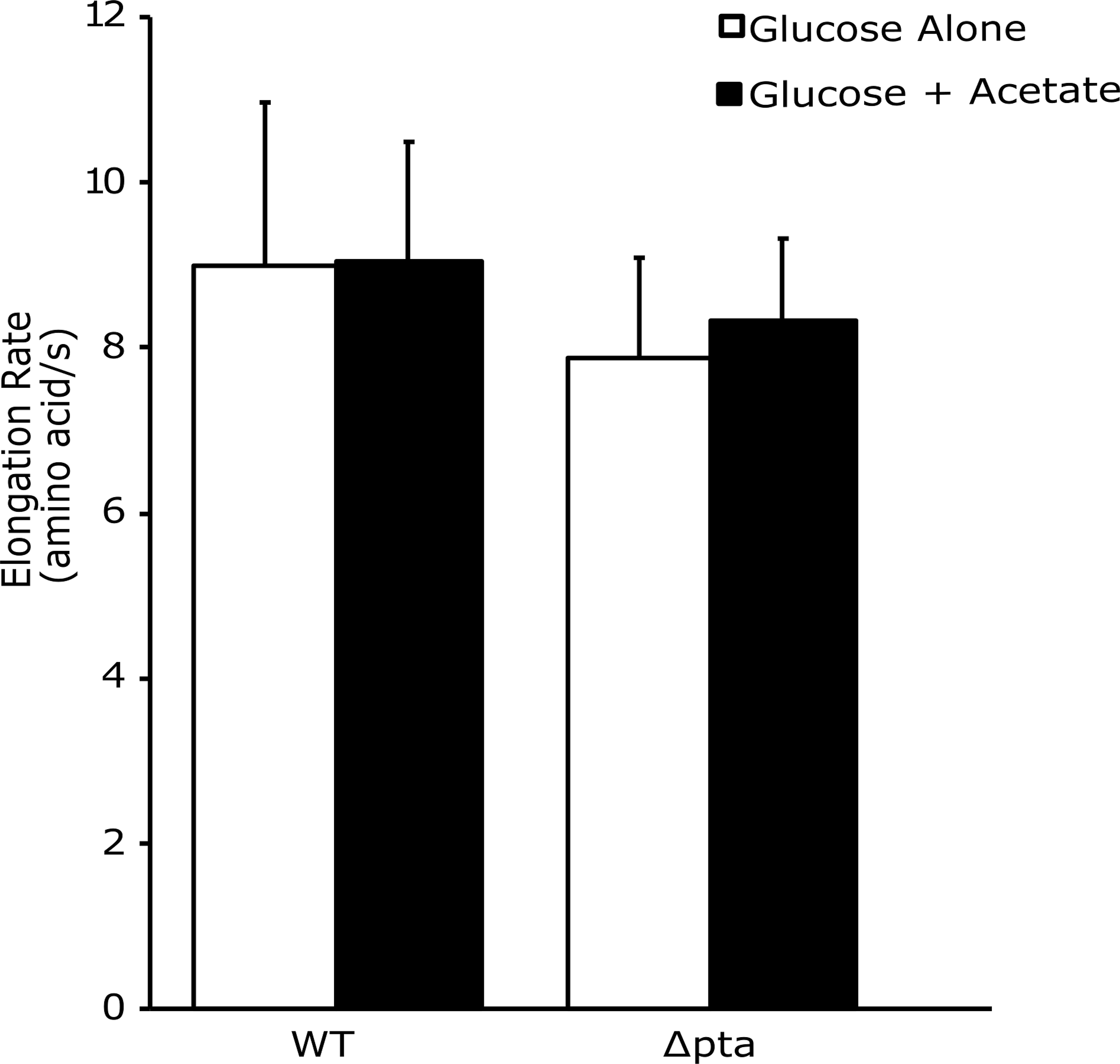
Conditions promoting acetylation do not affect elongation. Wild type MG1655 and an isogenic Δ*pt*a mutant were grown in MOPS + 0.2% glucose (white) or MOPS + 0.2% glucose supplemented with 0.27% acetate at 6 hours (black). β-galactosidase activity was induced at 8 hours and used to calculate the elongation rate in aa/s. Error bars represent the standard deviation from three replicates.

### High acetylation mutants promote ribosome dissociation as determined by polysome profiling

Our data suggest that acetyl donors, principally acetyl phosphate, inhibit translation in *E. coli*, most likely by acetylating ribosomal proteins. To gain further insight into the mechanism, we compared the polysome profiles for the wild type, a Δ*ackA* (high acetylation in a rich medium) mutant, and a Δ*pta* (low acetylation in a rich medium) mutant following 10 h of growth in TB7 containing 0.4% glucose (**Fig. 4**). A slight growth defect was observed in the Δ*ackA* and Δ*pta* mutants (**Fig. S2**) The wild-type profile exhibited a large peak associated with the 70S ribosome and smaller peaks associated with the 30S and 50S subunits. These peaks were verified using RNA gel electrophoresis (**Fig. 5A**). By contrast, the peak associated with the 70S ribosome was smaller in the Δ*ackA* (high acetylation) mutant, comparable in size with the peaks associated with the 30S and 50S subunits (**Fig. 4**). The profile for the Δ*pta* (low acetylation) mutant had larger 30S and 50S peaks and more polysomes compared to wild type but is more like wild type than the Δ*ackA* profile. As a control, we also profiled a complemented Δ*ackA* mutant (Δ*ackA* λ*att::ackA)* and Δ*pta* mutant (Δ*pta* λ*att::pta).* They exhibited polysome profiles identical to the wild type. These results suggest that conditions associated with high lysine acetylation favor dissociated subunits and that there is a more subtle effect associated with low acetylation.

**Figure 4.**
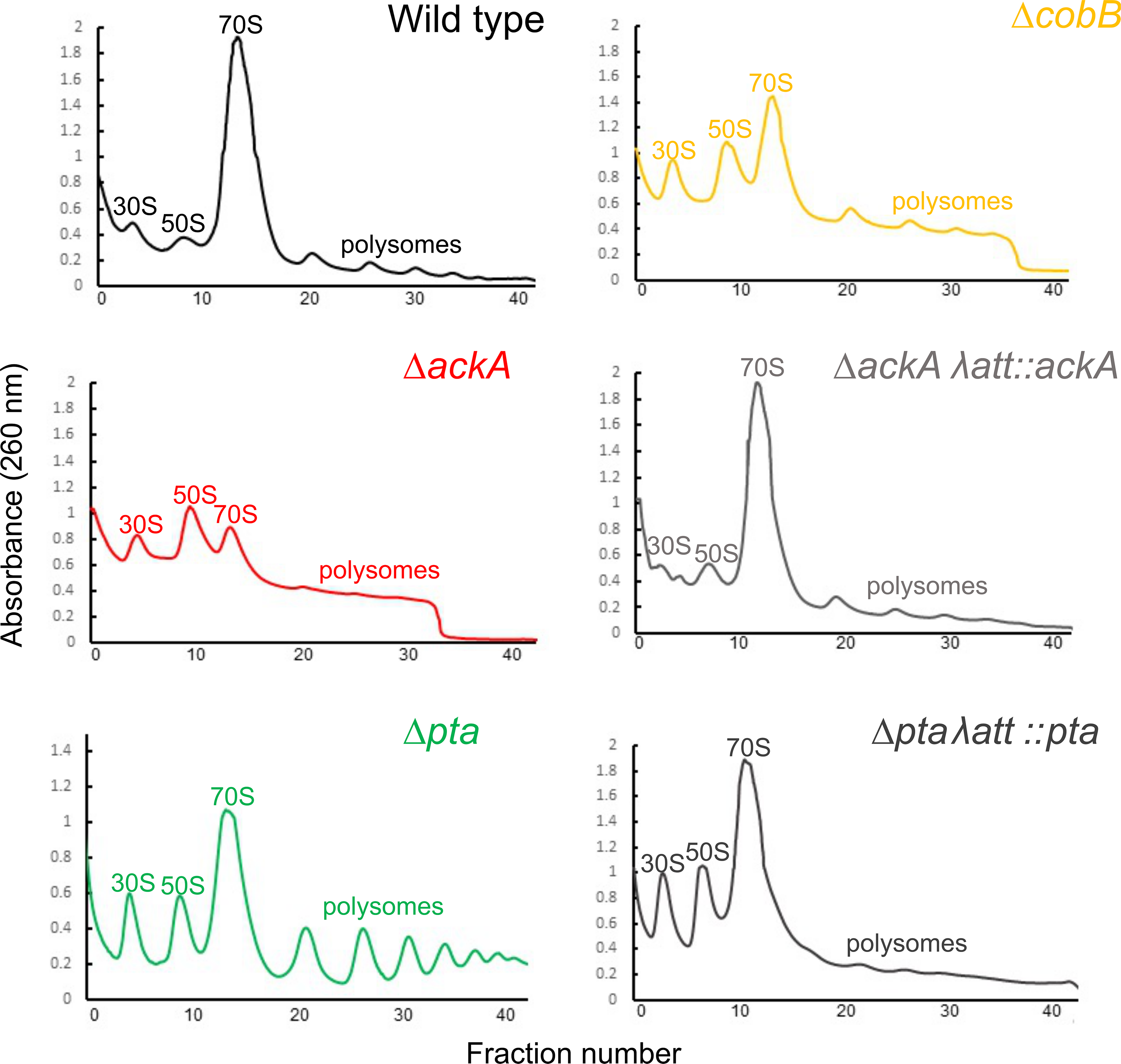
High acetylation mutants promote ribosome dissociation. Polysome profiles of wild type BW25113 and a series of isogenic mutants grown for 10 hours in TB7 + 0.4% glucose. 30S and 50S subunit peaks are marked, 70S monosome peak and polysome peaks are also marked. Identity of each peak was confirmed by RNA gel (Figure 5A **and data not shown**).

**Figure 5.**
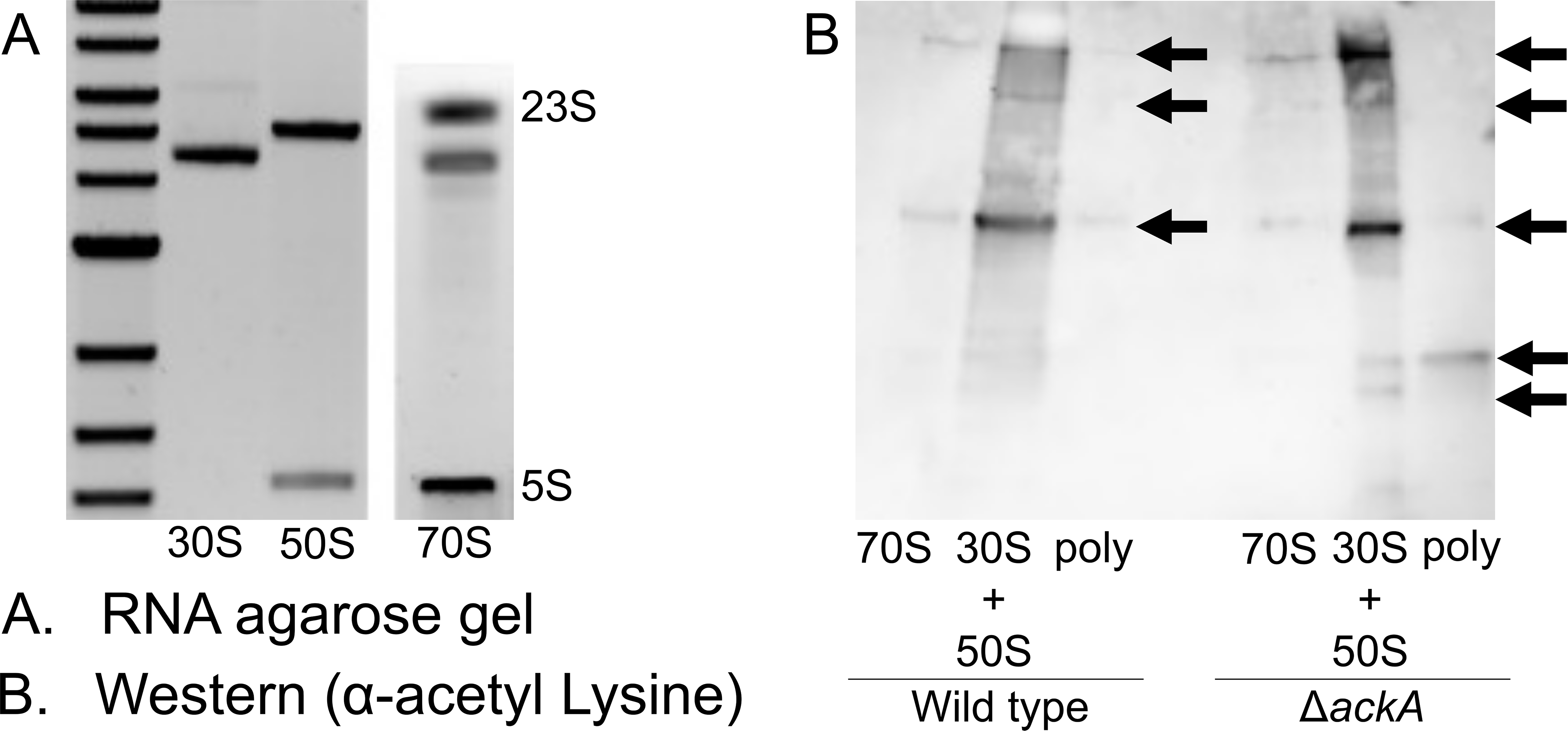
Analysis of polysomal gradient profiling fractions. (**A**) Agarose RNA gel for 30S, 50S, 70S and polysomal peak fractions collected from polysomal profile of wild type BW25113 grown for 10 hours in TB7 + 0.4% glucose. (**B**), Western blot using anti-acetylated lysine protein antibody for 30S +50S, 70S and polysomal peak fractions collected from polysomal profile of wild type BW25113 and its isogenic Δ*ackA* mutant grown for 10 hours in TB7 + 0.4% glucose. Loading was normalized to protein content. Arrows indicate bands of interest.

### Proteins associated with the 30S and 50S ribosomal subunits are more acetylated than those associated with the 70S ribosomal complex

The results discussed above demonstrate that mutants with highly acetylated proteomes have more disassociated ribosomes. This would suggest that the proteins within the dissociated 30S and 50S ribosomal subunits are more acetylated than those within the 70S ribosomal complex. To test this hypothesis, we performed Western blots using anti-acetyllysine antibodies on the pooled 30S and 50S, 70S, and polysome fractions from the wild type and Δ*ackA* mutant. For both strains, the 30S and 50S pooled fractions were more acetylated than the 70S or polysome fractions. This difference was more pronounced in the Δ*ackA* mutant, which also had a distinct band of acetylation in the polysome fraction not observed in the wild type. These results demonstrate that the disassociated 30S and 50S subunits contain more acetylated proteins than 70S complexes (**Fig. 5B**).

### Growth on acetate promotes ribosome dissociation

Because the growth conditions differed between our elongation and profiling results described above, we next performed polysome profiling for the wild type, the Δ*pta* mutant, and the Δ*ackA* mutant grown on 0.2% glucose versus 0.2% glucose supplemented with 0.27% sodium acetate (see **Fig. S1** for an example of cell growth under these conditions). In further support of a mechanism whereby acetylation promotes dissociated ribosomes, we observed a reduction in the peak associated with 70S ribosome and an increase in the peaks associated with the 30S and 50S subunits in both the wild type and Δ*pta* mutant when acetate was added to the growth medium (**Fig. 6**). The Δ*ackA* mutant, which is already acetylation high when grown in glucose, was less affected by the addition of acetate (**Fig. 6**). As growth on acetate is known to increase protein acetylation, these results further support the hypothesis that acetylation promotes ribosome dissociation (20). They also demonstrate that promotion of ribosome dissociation also occurs in the wild type and not just in high acetylation mutants.

**Figure 6.**
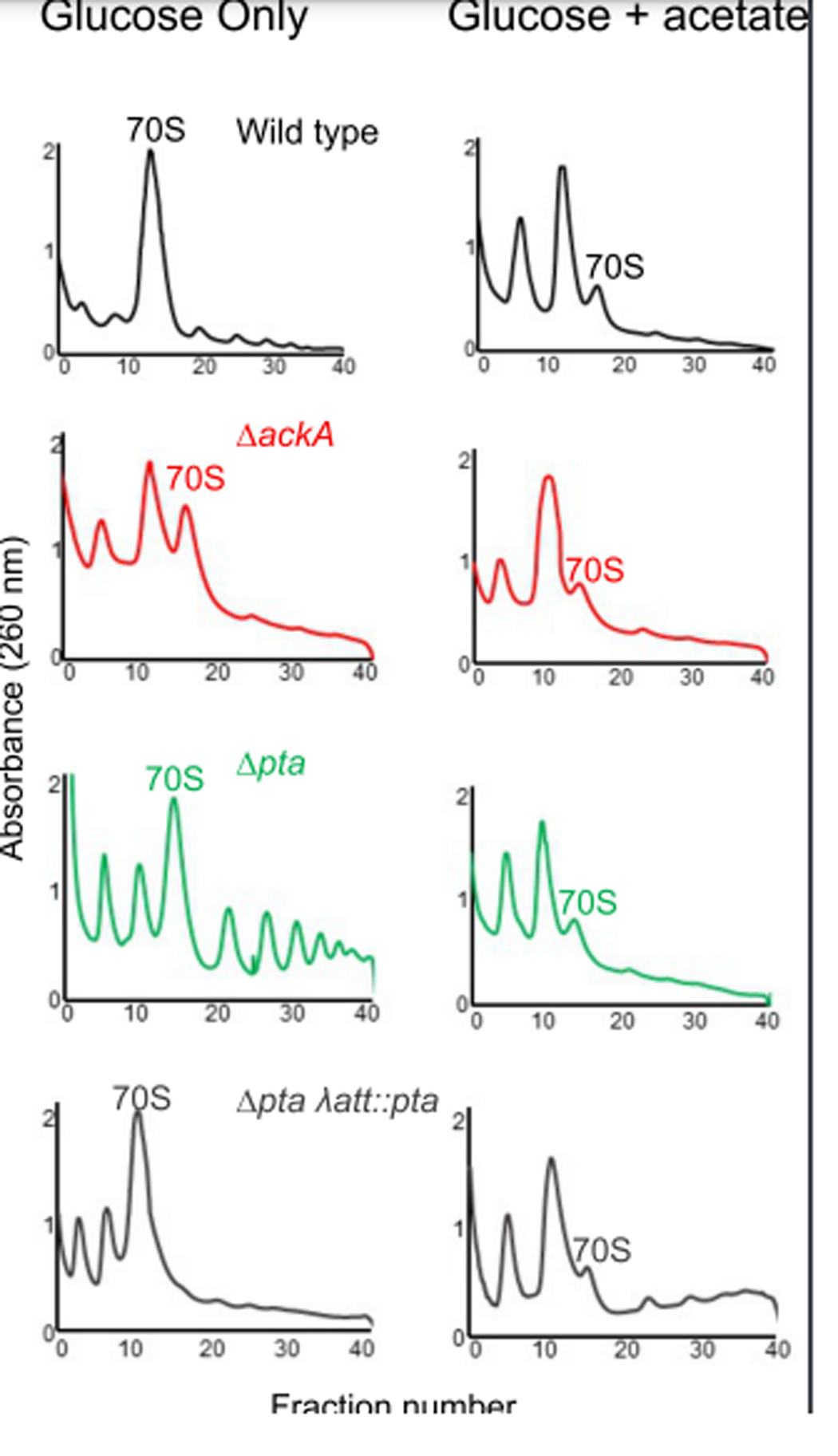
Growth on acetate promotes ribosome dissociation. Polysome profiles of wild type BW25113 and a series of isogenic mutants grown for a total of 10 hours in MOPS + 0.2% glucose or MOPS + 0.2% glucose supplemented with 0.27% acetate after 6 hours.

### Acetylation increases ribosome dissociation in wild type *E. coli* beginning in late exponential phase

Lysine acetylations accumulate in *E. coli* during the transition into stationary phase. Therefore, we hypothesized that we would not observe any significant differences in the polysome profiles for the wild type and Δ*ackA* mutant during exponential phase growth; rather, these differences would become significant only after entry into stationary phase. To test this hypothesis, we polysome profiled the wild type and Δ*ackA* mutant at multiple times along the growth curve (**Fig. 7**). While there was little difference in the profiles at early time points, the profiles for the Δ*ackA* mutants diverged from the wild type at later time points, as the cells exit exponential growth (**Fig S2**). In particular, the peak associated with the 70S ribosomal complex was reduced and the peaks for the 30S and 50S subunits increased. These differences persisted as the cells entered stationary phase.

**Figure 7.**
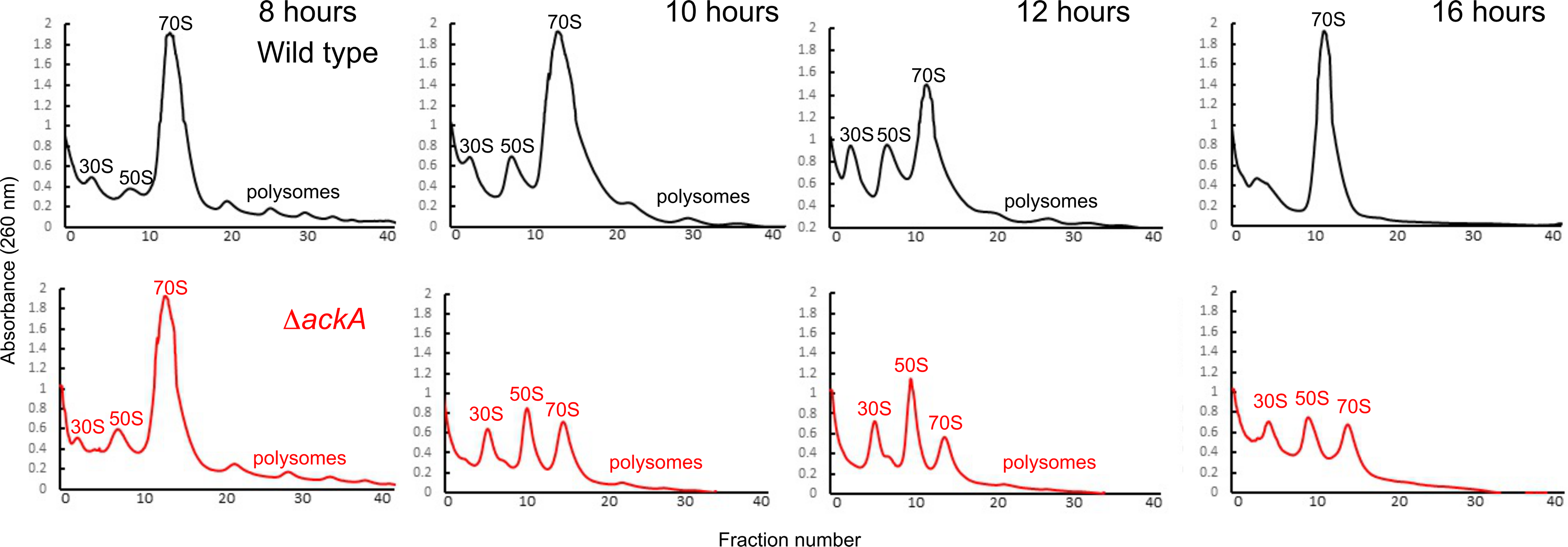
Polysomal gradient profiles for different time points. Polysome profiles of wild type BW25113 and an isogenic Δ*ackA* mutant grown in TB7 + 0.4% glucose over times noted. Percentage of ribosomes in 30S, 50S, and 70S fractions based on area under the curve calculations are provided in **Table 2**.

**Table 1.**
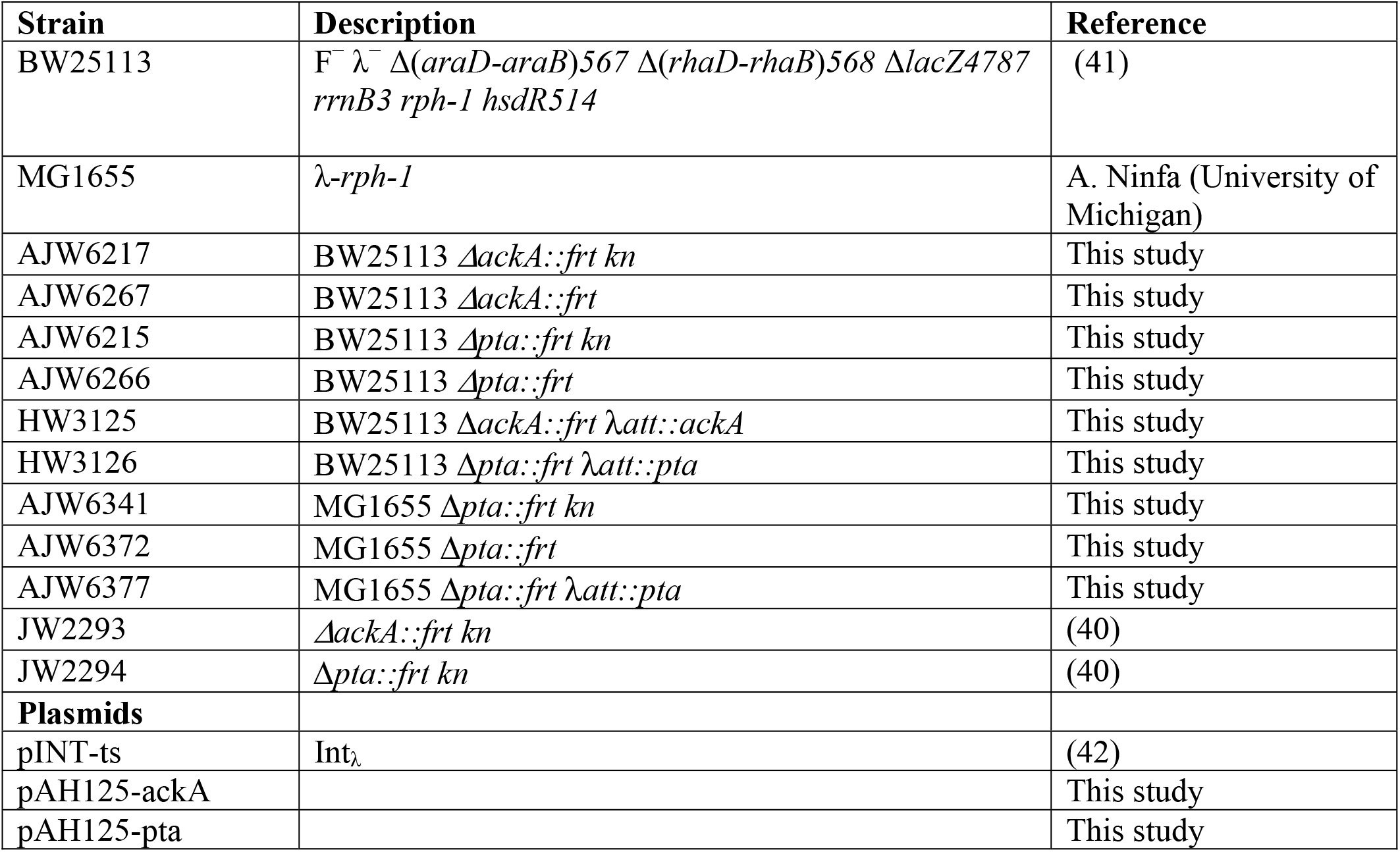
Bacterial strains and plasmids

**Table 2.**
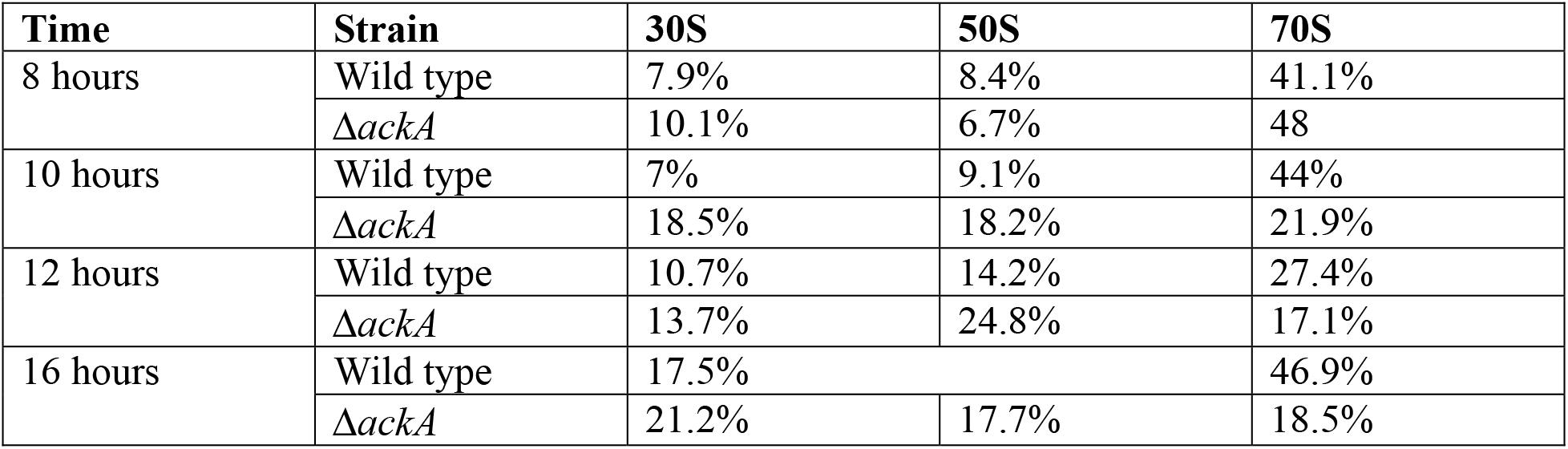
Portion of ribosomes in 30S, 50S, and 70S fractions over time

Interestingly, we observed increased ribosome dissociation when the wild-type cultures entered early stationary phase. Not unlike the Δ*ackA* mutant, we observed larger peaks associated with the 30S and 50S subunits. However, when wild-type cultures entered stationary phase, the peaks associated with the 30S and 50S subunits decreased, suggesting that dissociation was a transient phenomenon associated with the transition to stationary phase. This dynamic was not observed in the Δ*ackA* mutant. Whether transient dissociation is due to acetylation is not known, although Western blots indicated that dissociated ribosome subunits are more acetylated than the 70S complex.

## DISCUSSION

Ribosomal proteins are highly acetylated in diverse species of bacteria. A recent study suggested that acetylation of ribosomal proteins inhibits translation by reducing the rate of elongation (24). While our data suggests a more complex mechanism involving ribosome association/disassociation instead of elongation, both the previous and present studies demonstrate that acetylation affects translation. In our work, this was most clearly established when we measured protein production using a cell-free transcription/translation system. These experiments found that the addition of the acetyl donors, acetyl phosphate or acetyl-CoA, inhibits translation but not transcription in a dose-dependent manner. Our profiling experiments further demonstrate that fewer ribosomes form 70S complexes under conditions promoting high protein acetylation, either in mutants or by adding acetate to the growth medium. Moreover, the dissociated 30S and 50S subunits are more acetylated than the 70S complex, both in the wild type and the high acetylation *ΔackA* mutant. Taken together, these results demonstrate that ribosomal protein acetylation inhibits the ability of the ribosome subunits to form the 70S complex. Interestingly, we did not see any effect on translation elongation, which did not vary under condition of high and low acetylation.

While the exact mechanism remains opaque, the simplest explanation is that acetylation inhibits translation initiation, where the 30S and 50S subunits associate with initiation factors, tRNA, and mRNA to form the 70S complex. Ribosomal lysine residues are important for subunit association, with four of the eight inter-subunit bridges containing a lysine residue in *E. coli*. Among these interface residues, however, only K36 on the ribosomal protein L19 is known to be acetylated in *E. coli* (13, 17, 28). Other promising acetylation sites include three conserved lysine residues (K81, K84, and K100) on ribosomal protein L12, also known to be acetylated in *E. coli*. These conserved residues are located on a helix known to bind IF2, EF-Tu, EF-G, and RF3 (6, 29). IF2 is important for rapid subunit association during initiation (30–32). Mutations to K81 and K84 drastically impair subunit association and altering the complementary charges between K84 of L12 and D508 of IF2 impairs ribosomal subunit association (33). Because acetylation neutralizes the positive charge of lysine residues, it likely inhibits association by disrupting salt bridges along the subunit interfaces. While this mechanism is still speculative and requires further investigation to validate, it nonetheless provides one plausible explanation for how acetylation inhibits ribosomal assembly.

Proteins are most acetylated when the culture enters stationary phase (10, 13, 18). We observed a similar dynamic in our polysome profiling experiments. Indeed, subunit dissociation was most pronounced when cells were harvested from late exponential/early stationary phase cultures, both in the wild type and high acetylation Δ*ackA* mutant. One key difference is that disassociation occurred earlier in the Δ*ackA* mutant than the wild type and, for the *ackA* mutant, continued throughout stationary phase. These results are expected, as the Δ*ackA* mutant accumulates acetyl phosphate due to the loss of acetate kinase. In other words, we expect that proteins are more acetylated in the Δ*ackA* mutant even during exponential phase growth, as shown in prior work (12, 18). The other key difference is that inhibition of ribosome association is transient in the wild type: subunit levels peaked in early stationary phase but then disappeared as the cultures fully entered stationary phase. In the Δ*ackA* mutant, by contrast, dissociated subunits are observable well into stationary phase. Whether these latter differences are solely due to acetylation is not known, but the results clearly demonstrate that translation regulation by acetylation is a dynamic phenomenon that is growth phase specific.

*E. coli* produces acetate when the growth rate exceeds the respiratory capacity of the cell (20). When this occurs, the cells divert excess carbon flux towards acetate production. This enables cells to capture some energy that would otherwise be lost due to the inability to completely oxidize sugars at high rates. Acetate is also produced when carbon cannot be used to make biomass. This occurs when other essential nutrients/elements are depleted from the environment. For example, when nitrogen is depleted, the cells are unable to produce amino acids and instead divert excess carbon to acetate (34, 35). Under these conditions, protein acetylation is high (10).

Protein acetylation is tightly coupled to acetate metabolism in *E. coli*(*7*). This would suggest that acetylation of ribosomal proteins enables cells to couple translation and metabolism using a mechanism distinct from the stringent response. When a cell has a reduced need for protein synthesis, it has little need for ribosomes. This problem is partially addressed by the stringent response, where uncharged tRNAs induce the production of ppGpp(p) (36, 37). This, in turn, decreases transcription of the ribosomal operons. However, this mechanism only explains the rate of ribosome production. Because the cells cease to grow, existing ribosomes will not be “diluted away.” The cell, in other words, needs some mechanism for shutting down translation from pre-existing ribosomes. Our hypothesis is that this is achieved, at least in part, through acetylation. Such a mechanism would explain why acetylation inhibits translation. Whether this mechanism is coupled with ribosome hibernation and the formation of the inactive 100S complex is presently unknown.

We conclude by addressing the question of elongation. As noted above, a previous study found that acetylation inhibits translation by reducing the rate of elongation (24). In the present study, we did not observe any significant changes in the elongation rate. Rather, our profiling experiments strongly suggest that acetylation inhibits the association of the 30S and 50S ribosomal subunits. One likely explanation for these discrepancies is that different growth conditions were employed. In our experiments, we grew wild type and Δ*pta* mutant cells in minimal media containing glucose in the presence or absence of acetate. The previous study compared elongation in the wild type and a Δ*pta* mutant during growth in minimal medium with acetate as the sole carbon source. In our experience, Δ*pta* mutants grow much more slowly than the wild type when acetate is the sole carbon source (20, 38). Whether decreased growth is due to the lack of acetylation or a byproduct of growing a mutant defective in acetate metabolism on acetate is not known. Therefore, we supplemented the growth medium with glucose so that we could compare elongation in the wild type and Δ*pta* mutant in the presence or absence of acetate. Further work will be necessary to resolve these discrepancies.

## MATERIALS AND METHODS

### Bacterial strains, media, and growth conditions

All strains used in this work are derivatives of *E. coli* K-12 strains BW25113 or MG1655. Mutants were constructed by generalized transduction with P1kc, as described previously (39), with the Keio collection providing the appropriate deletions (40) with the exception of the Δ*ackA* λ*att::ackA* and the Δ*pta::frt* λ*att::pta* complementation mutants. First, the *ackA* or *pta* gene was deleted using the method of Dansenko and Wanner (41). Then, Gibson assembly was used for plasmid construction to ligate the *ackA* gene (MG1655 genomic region 2411492-2412445) and the *pta* gene (MG1655 genomic region 2412769-2414943) containing their respective promoter regions with the pAH125 vector, according to manufacturer’s instructions. The constructed plasmids were integrated into the λ attachment site in the chromosome using CRIM method with pINT-ts helper plasmid (42).

Cells were cultured overnight in 5 ml lysogeny broth (LB) and subcultured in 250 mL flasks containing 50 mL TB7 (10 g/liter tryptone buffered at pH 7.0 with 100 mM potassium phosphate), TB7 supplemented with 0.4% glucose, or MOPS minimal medium with 0.2% glucose as a carbon source for the times noted. When noted, MOPS cultures were supplemented with 0.27% acetate at 6 hours. All cultures were grown at 37 °C and aerated at 225 rpm with a flask-to-medium ratio of 5:1.

### Cell-free transcription/translation assay

All experiments were performed using the myTXTL Sigma 70 Master Mix Kit and P70a(2)-deGFP positive control plasmid (Arbor Biosciences). Briefly, 15 μL reactions were prepared by combining 12 μL of master mix and plasmid and 3 μL acetyl phosphate or acetyl-CoA at desired concentration. Distilled H_2_O was used for the control. Reactions were incubated for 2 hours at 37 °C in heat block. Reactions were stopped on ice and fluorescence measured. A standard GFP curve was used to calculate the amount of GFP synthesized.

### Quantitative PCR

RNA was isolated from cell-free reactions using the MasterPure Complete DNA and RNA Isolation kit (Epicenter). After RNA isolation, cDNA was prepared using the iScript cDNA Synthesis kit (BioRad). A standard curve for qRT-PCR was prepared using *E. coli* B gDNA, iTaq Universal 2x SYBR green (BioRad) and 16S primers (Forward primer: CGGTGGAGCATGTGGTTTA, Reverse primer: GAAAACTTCCGTGGATGTCAAGA). Samples, no template controls, and no iScript controls were combined with iTaq Universal 2x SYBR green (BioRad)) and primers for *deGFP* (Forward primer: GCACAAGCTGGAGTACAACTA, Reverse primer: TGTTGTGGCGGATCTTGAA). Reactions were carried out using the CFX Opus 96 Real-Time PCR System (BioRad). Expression of *deGFP* was calculated relative to the no acetyl phosphate control.

### Elongation rate assay

Translation elongation rates were measured using the LacZ induction assay (21, 27). Briefly, strains were grown overnight in MOPS + 0.2% glucose and sub-cultured to an OD_600_ 0.1 in 50 mL of MOPS + 0.2% glucose. Cultures were incubated shaking at 37 °C until stationary phase. Acetate cultures were supplemented with 0.27% acetate at 6 hours. At stationary phase, cultures were induced with 5 mM isopropyl β-D-thiogalactopyranoside (IPTG). Upon induction, at 30-s intervals, 100 μL of culture was harvested into pre-chilled Eppendorf tubes containing 5 μL chloramphenicol (34 mg/mL) for 10 time points. Samples were snap frozen and stored at -80 °C prior to LacZ assay. The assay was largely adapted from the traditional Miller’s colorimetric method but utilized the fluorescent substrate 4 methylumbelliferyl-D-galactopyranoside (MUG) instead of O-nitrophenyl-β-D- galactopyranoside (ONPG)(43, 44). Thawed samples were incubated with 400 μL Z-Buffer for 10 minutes at 37 °C before 50 μL MUG (2 mg/mL) was added to each sample. Reactions were incubated at 37 °C for 30 minutes then stopped with 250 μL 1 M sodium carbonate. Fluorescence was measured in black-sided 96-well plates (excitation 360 nm, emission 460 nm). LacZ inductions curves were made by plotting LacZ activity on the y-axis and time post- induction on the x-axis and further analyzed by square root plot to obtain the lag time for first LacZ molecule synthesis (T_first_)(45). LacZ is 1024 amino acids in length, and the translation elongation rate is calculated as 1024/T_first_.

### Polysome profiles

*E. coli* cultures were grown overnight in LB and subcultured the next morning in 50 mL of TB7 + 0.4% glucose or MOPS minimal media + 0.2% glucose to an OD_600_ of 0.02. Cultures were then grown at 37 °C. When noted, cultures were supplement with 0.27% sodium acetate at 6 hours. At harvest time, 50 μL chloramphenicol (100 mg/mL) was added, and cultures were rapidly cooled and pelleted by centrifugation at 4,000 x *g* for 15 minutes at 4 °C. Cells were then lysed in a buffer consisting of 10 mM Tris-HCl (pH 8.0), 10 mM MgCl_2_, and 1 mg/mL lysozyme by three freeze-thaw cycles. After the final freeze-thaw, 15 μL 10% sodium deoxycholate was added and cellular debris was pelleted by centrifugation at 9,400 x *g* for 10 minutes at 4°C. The supernatant was collected and stored at -20 °C for profiling.

Profiles were run on a 10%-40% sucrose gradient prepared using a sucrose buffer consisting of 20 mM Tris-HCl (pH 7.8), 10 mM MgCl_2_, 100 mM NH_4_Cl, and 2 mM DTT. Gradients were prepared using (Biocomp Gradient Master 108). Each gradient was loaded with 300 μL of *E. coli* lysate and spun using an SW-41 rotor in an ultracentrifuge at 175,117 x *g* for 3 h 45 min at 4 °C. Gradients were fractionated using the ISCO/Brandel fractionation system by injecting a 50% sucrose solution below the gradient at 1.5 mL/min. Ribosomes were detected by the system’s UV spectrophotometer at 254 nm. Fractions were stored at -20 °C for future analysis by Western blotting.

### RNA purification and electrophoresis

Ribosome peak fractions were pooled. The 30S and 50S ribosomal peaks were processed directly but the 70S ribosome fraction and polysome fractions were dissociated into 50S and 30S subunits and bound mRNA was removed by loading each fraction onto a 10%-45% sucrose gradient made in disassociation buffer (20 mM HEPES pH 7.5, 5 mM β-mercaptoethanol, 5 mM MgCl_2_, 50 mM NH_4_Cl, 0.1 mM PMSF). The gradient was centrifuged as before, using an SW-41 rotor in an ultracentrifuge at 175,117 x *g* for 3 h 45 min at 4 °C. Gradients were fractionated using the ISCO/Brandel fractionation system. To all individual pooled ribosomal fractions 1.5× volume of TRIzol reagent (Invitrogen) was added. Sample tubes were shaken for 15 s and incubated at room temperature for 10 min. Samples were then layered onto a Direct-zol RNA MiniPrep (Zymo Research) spin column, and RNA was extracted according to manufactures directions. Polysomal RNA can be isolated using TRIzol alone, but we were able to obtain cleaner RNA using the Direct-zol RNA MiniPrep (Zymo Research) in addition to TRIzol (Invitrogen). To visualize RNA, mix 0.5 ug of the purified RNA, with 1.5x volume of deionized formaldehyde, and RNA loading buffer (0.25% bromophenol blue, 0.25% xylene cyanol, 30% glycerol) and load onto a 1.2% agarose gel made in TBE (45 mM Tris-borate, 1 mM EDTA pH 8.0) using diethyl pyrocarbonate (DEPC) treated water. The gel was run for 45 minutes at 100 Volts and RNA bands were visualized using SYBR Green II RNA Gel Stain (ThermoFisher Scientific).

### Anti-acetyllysine Western blots

The protein concentrations within the fractionated sample loaded onto the gel was normalized by total protein content using the bicinchoninic acid (BCA) assay (Thermo Scientific Pierce, Waltham, MA). Proteins were separated by 12% sodium dodecyl sulfate-polyacrylamide gel electrophoresis (SDS-PAGE). Gels were rinsed in transfer buffer (25 mM Tris, 192 mM glycine, 10% Methanol) and the proteins transferred onto a nitrocellulose membrane in transfer buffer for 1.5 hours at 100 V at 4 °C. After transfer, membranes were blocked with 5% milk in PBST (137 mM NaCl, 2.7 mM KCl, 10 mM Na_2_HPO_4_, 1.8 mM KH_2_PO_4_, 0.1% Tween) for 1 hour and washed with PBST four times for 5 minutes each. Primary rabbit anti-acetyllysine antibody (Cell Signaling, Danvers, MA) was diluted 1000-fold in 5% BSA, added to the membranes, and incubated in the cold room with shaking. The membrane was washed 4x with PBST for 5 minutes each and incubated for 1 hour in the dark at room temperature with anti-rabbit IgG HRP-linked secondary antibody (Cell Signaling, Danvers, MA) diluted 2000-fold in 5% milk. The membrane was washed 4x with PBST for 5 minutes each, incubated in ECL blotting substrate (Abcam) and imaged in the Protein Simple machine (Bio-Techne) (13, 19).

## Data availability

Data available upon request.

**Supplementary Figure 1. Growth in MOPS + 0.2% glucose with and without acetate supplementation over time.** Wild type MG1655 and an isogenic Δ*pt*a mutant were grown in MOPS + 0.2% glucose or MOPS + 0.2% glucose supplemented with 0.27% acetate at 6 hours (indicated by arrow). Optical density was measured at 600 nm. Each time point is the average of 3 biological replicates with error bars representing the standard deviation.

**Supplementary Figure 2. Growth in TB7 + 0.4 glucose over time.** Wild type BW25113 and a series of isogenic mutants grown for 16 hours in TB7 + 0.4% glucose. Optical density was measured at 594 nm.

## ACKNOWLEDGMENTS

We wish to thank Professor Hong Jin and Dr. Melissa Alves for advice concerning the polysome profiling experiments. We also wish to thank Thomas Bank for helpful conversations throughout the study. HEW and CVR were funded by DOE Center for Advanced Bioenergy and Bioproducts Innovation (U.S. Department of Energy, Office of Science, Office of Biological and Environmental Research under Award Number DE-SC0018420). Any opinions, findings and conclusions or recommendations expressed in this publication are those of the author(s) and do not necessarily reflect the views of the U.S. Department of Energy.

